# Charge-perturbation dynamics — a new avenue towards in silico protein folding

**DOI:** 10.1101/597039

**Authors:** Purbaj Pant, Ravi José Tristão Ramos, Crina-Maria Ionescu, Jaroslav Koča

## Abstract

Molecular dynamics (MD) has greatly contributed to understanding and predicting the way proteins fold. However, the time-scale and complexity of folding are not accessible via classical MD. Furthermore, efficient folding pipelines involving enhanced MD techniques are not routinely accessible. We aimed to determine whether perturbing the electrostatic component of the MD force field can help expedite folding simulations. We developed charge-perturbation dynamics (CPD), an MD-based simulation approach that involves periodically perturbing the atomic charges to values non-native to the MD force field. CPD obtains suitable sampling via multiple iterations in which a classical MD segment (with native charges) is followed by a very short segment of perturbed MD (using the same force field and conditions, but with non-native charges); subsequently, partially folded intermediates are refined via a longer segment of classical MD. Among the partially folded structures from low-energy regions of the free-energy landscape sampled, the lowest-energy conformer with high root-mean-square deviation to the starting structure and low radius of gyration is defined as the folded structure. Upon benchmark testing, we found that medium-length peptides such as an alanine-based pentadecapeptide, an amyloid-β peptide, and the tryptophan-cage mini-protein can fold starting from their extended linear structure in under 45 ns of CPD (total simulation time), versus over 100 ns of classical MD. CPD not only achieved folding close to the desired conformation but also sampled key intermediates along the folding pathway without prior knowledge of the folding mechanism or final folded structure. Our findings confirmed that perturbing the electrostatic component of the classical MD force field can help expedite folding simulations without changing the MD algorithm or using expensive computing architectures. CPD can be employed to probe the folding dynamics of known, putative, or planned peptides, as well as to improve sampling in more advanced simulations or to guide further experiments.

**Author summary:** Folding represents the process by which proteins assemble into biologically active conformations. While computational techniques such as molecular dynamics (MD) have provided invaluable insight into protein folding, efficient folding pipelines are not routinely accessible. In MD, the behavior of the studied molecule is simulated under the concerted action of multiple forces described by mathematical functions employing optimized parameters. Using non-native parameters effectively perturbs the MD force field. We show that this can be exploited to help expedite folding simulations. Specifically, we developed charge-perturbation dynamics (CPD), an MD-based simulation approach that involves periodically perturbing the force field by using non-native atomic charges. For folding medium-length peptides such as the tryptophan-cage mini-protein starting from the extended linear structure, CPD is much faster than other MD-based approaches while using the same software, hardware, and know-how required for running classical MD simulations. Furthermore, CPD not only achieves folding close to the desired conformation but also samples key intermediates along the folding pathway without prior knowledge of the folding mechanism or final folded structure. CPD can be employed to probe the folding dynamics of known, putative, or planned peptides, as well as to generate different conformations that can guide further experiments or more advanced simulations.

## Introduction

Folding represents a complex phenomenon by which proteins assemble into biologically active conformations. Misfolding events can have detrimental and sometimes catastrophic effects on the ability of the protein to perform its function, on its distribution in various cell compartments, and on its recognition by other species^1^. Therefore, the study of protein folding, unfolding, and misfolding is critical to our understanding and manipulation of pathophysiological mechanisms and biotechnological processes^2-4^. Great advances in the field of in silico modelling, including molecular dynamics (MD), have helped understand important aspects of folding, such as the fact that protein-folding energy landscapes are funnel-shaped, or that proteins fold in units of secondary structures^5-7^.

MD, which simulates the behavior of a molecular system under the resultant action of a set of forces (i.e., force field), is a very popular computational approach to study protein dynamics, as it can provide information about folding and unfolding pathways, intra-protein interactions, intermediate and final structures, and timeline of folding events; however, the time-scale and complexity of folding are often not accessible via classical MD^8,9^. To address such limitations, enhanced MD techniques typically employ one or more of the following strategies: simplifying the representation of the protein structure^10^, constraining or restraining the simulation^11^, steering the simulation in a pre-specified direction^12^, enhancing the sampling of molecular conformations^13^, describing physical interactions more accurately^14,15^, and employing software and hardware platforms specifically dedicated to increasing the computational efficiency of MD calculations^16^. MD platforms that can achieve millisecond timescales^17,18^ are not routinely accessible, and thus most users still rely on local computational clusters, which typically do not have enough resources for efficiently folding small proteins. Moreover, the practical applicability of enhanced MD techniques is often limited to certain use cases (e.g., requiring a priori knowledge of the folded structure or of states along the folding pathway)^8,19^. Finally, setting up MD calculations that can make efficient use of such enhancement techniques represents a complex task, requiring advanced knowledge of modelling techniques.

In an effort to improve the availability of MD for investigations that involve protein folding or may benefit from information related to folding, we aimed to determine whether perturbing the electrostatic component of the classical MD force field can help promote folding on shorter time scales while using the same software, hardware, and know-how required for running classical MD simulations. To test this hypothesis, we developed an MD technique that relies on classical MD but includes short segments where the electrostatic component of the force field is heavily perturbed. We refer to this technique as charge-perturbation dynamics (CPD), although the perturbed component is modelled in the same way as the classical electrostatic component. We here describe the main principles of CPD and the results of a benchmark test for the folding of medium-length peptides. We compare the speed and accuracy of CPD with those of classical MD, and further compare the CPD folding results with those from independent studies that apply more complex methods and more expensive computational resources to fold the same peptides. While peptide folding simulations typically require at least hundreds of ns, we show that medium-length peptides can fold via CPD starting from their extended linear structure in under 45 ns (total simulation time), without any prior knowledge of the folding mechanism or final folded structure, and using the same software, hardware, and know-how required for running classical MD simulations.

## Results

### Principles of CPD

MD simulations describe the behavior of molecular systems under the influence of the force field. Various force fields have been proposed to date^20,21^. Commonly used force fields use approximate functions to describe the contributions of bonded interactions (e.g., torsion) and non-bonded interactions (e.g., electrostatic). Point charges residing at the position of each atom (i.e., atomic charges) are commonly used as parameters in the calculation of the electrostatic component. Because the concept of atomic charge represents a very crude approximation of electron density, there is no universally accepted quantitative definition of atomic charge. Current approaches for the calculation of atomic charges partition the molecular electron density obtained from quantum mechanical calculations (e.g., see Lee et al.^22^). However, such quantum mechanics-based approaches are rarely used in MD; instead, empirical, force field-specific approaches are typically applied, as they provide both speed and adaptability for the force field^23^. In fact, force-field parameters including atomic charges are optimized in an inter-dependent manner in a process known as force-field parameterization^24,25^, so that the overall force field may provide a reasonable description of the studied system (e.g., density at room temperature). Using non-native or even suboptimal parameters effectively perturbs the MD force field. We show that this characteristic can be exploited to help expedite folding simulations. The concept underlying CPD is that substantial conformational changes can be promoted during the MD simulation by perturbing the electrostatic component of the force field over brief segments of the simulation. Within CPD, the electrostatic component is perturbed by replacing the atomic charges native to the force field (i.e., the type of atomic charges used during force field parameterization) with non-native values. Specifically, the classical MD simulation (i.e., the MD simulation using native charges) is periodically intercalated with brief segments where non-native charges are used. The classical MD segments optimize the interactions within the secondary structure elements, while the perturbed MD segments allow these elements to reposition themselves relative to one another. This approach ultimately facilitates folding over a much shorter time scale, and thus requires significantly fewer computational resources than those necessary for folding using currently available techniques.

### Benchmarking a CPD protocol

To examine whether CPD can help expedite folding simulations, we implemented the above-described principles into a simple pipeline covering 45 ns of simulation time (Fig 1). The CPD protocol used in this study had two main stages: one focused on obtaining suitable sampling, and one focused on identifying and refining the folded structure. The detailed description of the pipeline is as follows, with the values in parentheses representing the exact settings used in the benchmark. Within stage I, the starting structure (a linear chain of amino acid residues) is minimized and equilibrated (1 ns). Subsequently, a segment of classical MD is run (400 ps), with snapshots taken at short intervals (4 ps). Thereafter, a brief segment (100 ps) of perturbed MD is run, during which the atomic charges native to the force field are replaced by non-native charges (conformation-dependent atomic charges obtained via an electronegativity equalization method)^26,27^. At the end of the perturbed MD segment, the secondary structure content is estimated for each snapshot recorded during this short segment, and the snapshot with the highest content of secondary structure is then used as the first frame of the subsequent classical MD segment. This sequence of steps consisting of a segment of classical MD, a segment of perturbed MD, and evaluation of secondary structure is iterated several times (50 times), giving a total simulation time of several tens of ns (25 ns) for stage I. Stage II follows, wherein a free-energy landscape is plotted based on two key measures calculated from each snapshot sampled in stage I, namely the root mean square deviation (RMSD) to the starting structure, and the radius of gyration (Rg). Since the starting structure was linear, the free-energy landscape built in this manner allows to identify partially folded structures in the low-energy regions, as such conformers are expected to have high RMSD and low Rg as indicators of folding^28^. Once the stage I snapshot with the lowest energy is identified, it serves as the initial structure for a longer segment of classical MD (20 ns), which represents stage II, with snapshots recorded at short intervals (4 ps). Finally, the overall free-energy landscape is built based on RMSD and Rg obtained from all snapshots (i.e., stage I and stage II). The lowest energy structure is provided as the final, folded structure. A detailed pseudocode of this protocol is available in the S1 Appendix and describes the setting up, running, and evaluating the results of the MD simulations.

**Fig 1.**
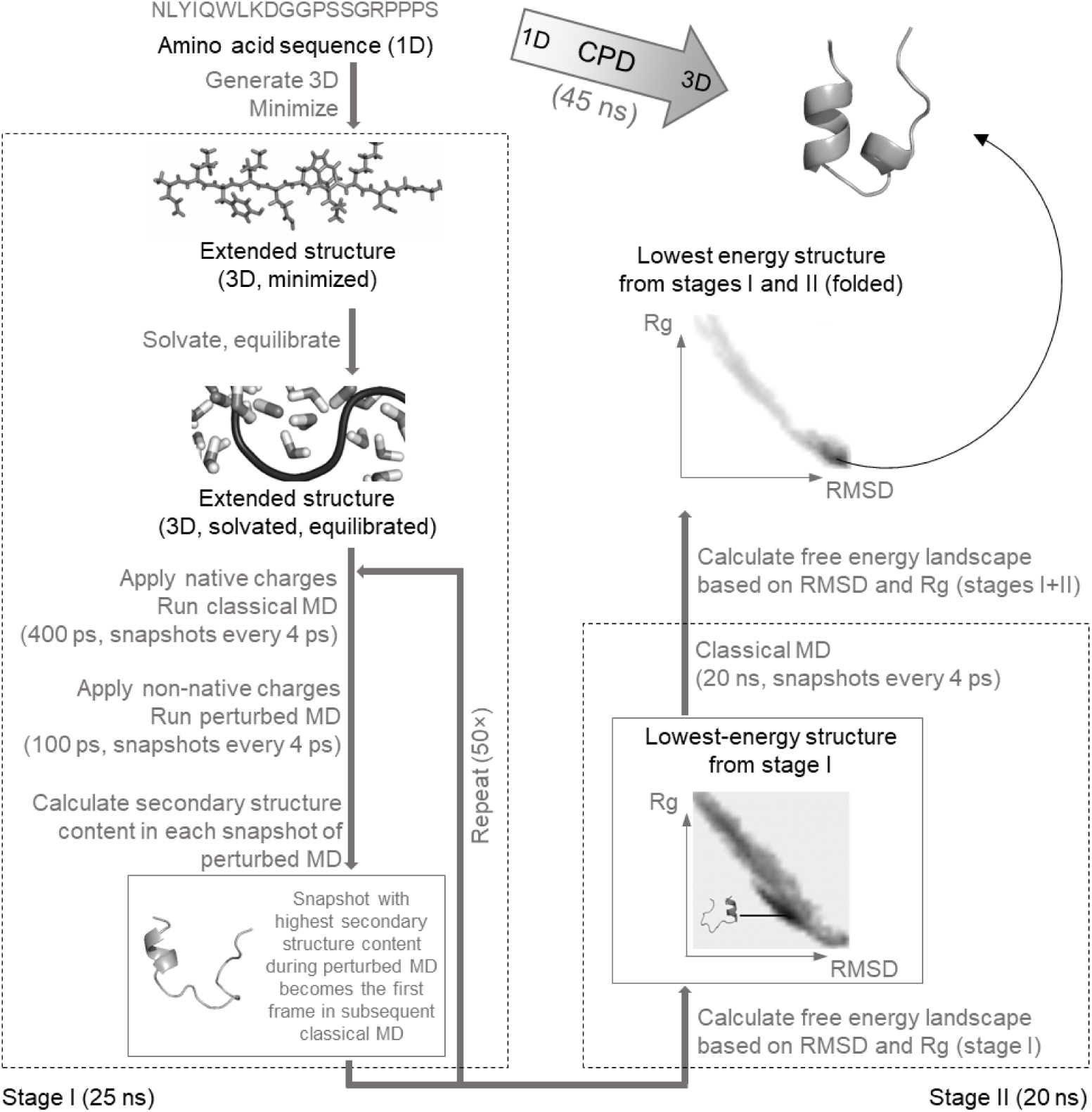
Schematic diagram of a charge-perturbation dynamics (CPD) protocol for expediting molecular dynamics (MD)-based folding simulations. CPD integrates segments of classical molecular dynamics (MD) with segments of perturbed MD, where the atomic charges native to the force field are replaced with non-native charges, resulting in a perturbation of the electrostatic component of the force field. The information provided in parentheses refers to the settings used in the present benchmark. All simulations started from the amino acid sequence. Stage I, which is focused on obtaining suitable sampling, consists of an initial step of minimization, solvation, and equilibration of the extended linear structure, followed by a set of alternating segments of classical and perturbed MD. With the exception of the very first segment of classical MD, the initial structure for each segment of classical MD corresponds to the snapshot with the highest content of secondary structure sampled in the preceding segment of perturbed MD. Stage II is concerned with identifying and refining the partially folded structure sampled in stage I. A free-energy landscape is plotted based on indicators of folding computed from the snapshots recorded in stage I, namely the radius of gyration (Rg) and the root mean square deviation (RMSD) relative to the minimized linear structure. A snapshot with the lowest energy from a region with high RMSD and low Rg is used as the initial frame in a final segment of classical MD. The overall free-energy landscape is built using snapshots from stage I and stage II, and the lowest-energy structure is extracted as the final folded structure.

Such a CPD protocol totaling 45 ns of simulation time was benchmarked against classical MD totaling 100 ns in terms of the ability to fold medium-length peptides commonly used for benchmarking protein folding techniques. All simulations started from the extended linear structure. In each case, triplicate simulations were run with different starting velocities, to verify that rapid folding is not a random event. The best results are shown in Fig 2, while the complete results are provided in S1 Fig and S2 Fig for CPD and classical MD, respectively.

**Fig 2.**
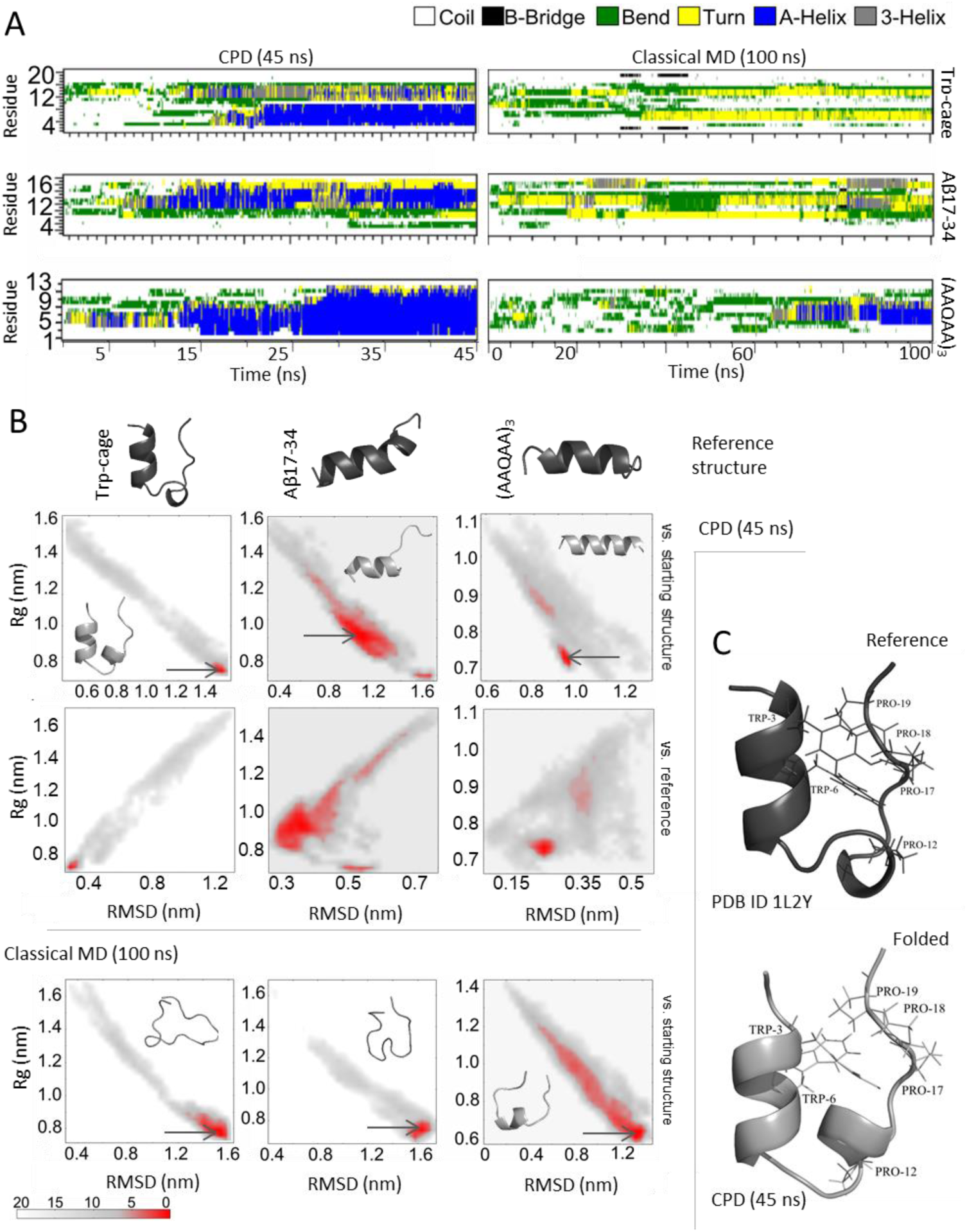
Benchmarking a charge-perturbation dynamics (CPD, 45 ns) protocol against classical molecular dynamics (MD, 100 ns) for folding medium-sized peptides. CPD, which consisted of two stages, achieved rapid folding of the tryptophan-cage mini-protein (Trp-cage), amyloid-β peptide Aβ17-34, and alanine-based pentadecapeptide (AAQAA)3. All simulations were run in triplicate, with different starting velocities for the atoms. Only the results of the best-performing runs are given here, whereas the complete results are available in S1 Fig and S2 Fig. (A) Formation of secondary-structure elements over the course of 45 ns of CPD (left column) and 100 ns of classical MD (right column). (B) Free-energy landscapes and lowest-energy structures obtained following 45 ns of CPD (top lane) and 100 ns of classical MD (bottom lane). The reference folded structures are also shown for comparison. The RMSD against the starting structure (linear) was used for detecting the lowest-energy structures. The RMSD against the reference structure is shown for comparison but was not used in during the simulation. (C) Detailed comparison of the hydrophobic core in the folded Trp-cage (reference, top; best CPD, bottom). Abbreviations: Rg, radius of gyration; RMSD, root mean square deviation.

### Folding the tryptophan cage (Trp-cage) in 45 ns of simulation time

The Trp-cage is an engineered mini-protein containing 20 amino acid residues (NLYIQWLKDGGPSSGRPPPS). The Trp-cage is often used in folding studies because it folds fast and because it is well studied both experimentally and theoretically. The Trp-cage consists of an N-terminal α-helix (residues 2–8), followed by a 3_10_-helix (residues 11– 14) (Fig 2B and 2C). Folding is cooperative and hydrophobically driven by the encapsulation of a Trp side-chain in a sheath of Pro rings (Fig 2C, top)^29^. Specifically, folding relies on the formation of a hydrophobic core in which Trp6 is buried in the center by residues Pro12, Pro17, Pro18, and Pro19^30^.

We subjected the linear extended structure of the Trp-cage to the CPD simulation protocol. Secondary structure elements formed already during stage I of the simulation, and the expected α-helix and 310-helix were stable throughout stage II (Fig 2A). In agreement with experimental findings, we found that the formation of the cage depends on the formation of the α-helix (i.e., the α-helix forms first), whereas the 3_10_-helix is less stable and likely to unfold before the rest of the structure under temperature stress^31^. The final folded structure (Fig 2B and 2C) was within 2.86 Å (backbone RMSD) of the reference folded Trp-cage (PDB ID 1L2Y)^29^. Since the pairwise RMSD values for reported NMR models of the Trpcage (i.e., those included in PDB ID 1L2Y) range from 0.54 Å to 1.39 Å, it can be considered that the differences between the reference folded structure and the structure folded via CPD are most likely related to force field-specific limitations^32^. In the CPD-folded structure, the hydrophobic core is present, with Trp6 at its center, packed by hydrophobic residues including Pro and Tyr (Fig 2C, bottom). The specific Trp6 interactions are similar to those reported in previous simulations^30^, but different than those in the reference folded structure reported based on NMR findings (Fig 2C, top)^29^. Nevertheless, mutational studies have shown that non-specific indole/backbone interactions might be more relevant for folding than specific indole-proline interactions^33^. In one of the CPD runs, the Trp3 side chain is not closely packed with the central Trp6, but fully exposed (see S1 Fig). This feature was previously reported in simulations using completely different setups than our own^32^. Furthermore, the final folded structure from the CPD run does not exhibit the salt-bridge between Asp9 and Arg16, which appears in the reference folded structure^29^. The formation of this salt bridge and its role in the stability of the Trp-cage structure are somewhat controversial. Some have speculated that this salt bridge enables fast folding and contributes to the stability of the folded structure^29,32,34^, while others claimed that the fully folded state can be obtained only after breaking of the salt bridge^30,35^, or that the salt bridge is not required at all for folding^36-38^. Thus, the CPD-folded structure of the Trp-cage is considered to be sufficiently close to the native state. By comparison, the classical MD simulation starting from the extended structure of the Trp-cage did not show formation of any significant helical element even after 100 ns (Fig 2A). The only interesting feature of the classical MD simulations was that the N-terminal residues occasionally assembled into a short-lived 3_10_-helix.

### Folding a water-soluble amyloid-β (Aβ) peptide in 45 ns of simulation time

The folding of the Aβ peptide and various Aβ segments has been the topic of intense study because of its association with the progression of Alzheimer’s disease^39^. The water soluble Aβ17-34 is an 18-residue (LVFFAEDVGSNKGAIIGL) segment of Aβ, which, in aqueous solutions under physiological conditions, was observed to adopt an α-helical structure for residues 19–26 and 28–33, with a kink around residues 26–28 (Fig 2B)^40^.

We subjected the linear extended structure of Aβ17-34 to the CPD simulation protocol. Already during stage I, we observed the formation of an α-helical element in the second half of Aβ17-34, along with the expected kink. Stage II sampled the full length of the α-helix between residues 28 and 33, and the kink at the expected position between residues 26 and 27 (Fig 2A). Interestingly, an intermediate with 3_10_-helical structure between residues 25 and 33 was sampled to a significant extent. This observation is in agreement with a report based on a completely different simulation setup, where a similar intermediate with 3_10_-helical structure between residues 26 and 28 was sampled in Aβ21-30^41^. The final CPD-folded structure does not exhibit a well-defined α-helical structure prior to the kink (residues 19–26) (Fig 2B) but is within 3.42 Å (backbone RMSD) of the reference folded structure of Aβ17-34 (PDB ID 2MJ1)^40^. It is worth noting here that the reference folded structures from the NMR ensemble differ from one another by up to 2.79 Å. Moreover, in the NMR experiment, the Aβ17-34 contained two additional glutamic acid residues at each terminus, which increased solubility and stabilized the helical structure in aqueous solution^40^; since our simulations only included residues 17–34 (i.e., without the terminal residues added to stabilize the helix), we consider that CPD achieved a satisfactory proportion of helical structure.

By comparison, the classical MD simulation starting from the extended structure of Aβ17-34 showed formation of significant helical elements only after 80 ns, but these were stable only for approximately 10 ns (Fig 2A). An interesting feature of the classical MD simulation was that the 3_10_-helical elements seemed to be more stable than the α-helical elements.

### Folding an alanine-based decapentapeptide in 45 ns of simulation time

Alanine-based peptides adopt significant populations of helical structures in aqueous solution^42^. The alanine-based decapentapeptide (AAQAA)3 was shown experimentally to exhibit significant helical content^43^. Successful simulations of the folding of (AAQAA)3 have relied on an accurate description of interactions^44^or employed enhanced sampling techniques^45,46^.

We subjected the linear extended structure of (AAQAA)3 to the CPD simulation protocol. Helical segments formed already during stage I and were amply sampled in stage II (Fig 2A). Since no experimental structure has been published for (AAQAA)3 to date, we used as reference the latest folded structure reported for (AAQAA)3^47^. The final CPD-folded structure was within 2.53 Å (backbone RMSD) of the reference. Interestingly, stage II also sampled 3_10_-helical elements, in agreement with the suggestion that such elements appear as intermediates during folding^48,49^.

By comparison, the classical MD simulation starting from the extended structure of (AAQAA)3 showed formation of a stable α-helical element in the central part of the peptide only after 70 ns. However, the folding did not progress significantly during the following (and final) 30 ns.

## Discussion

The aim of the current study was to determine whether perturbing the electrostatic component of the MD force field can help expedite MD-based folding simulations. We proposed and successfully validated a simple CPD protocol for rapidly folding medium-sized peptides. We further discuss key aspects of the CPD framework.

### Role of charge perturbation

In principle, any charge calculation scheme can be used for generating the non-native charges used in the perturbed MD segments, even if the representation of electrostatic interactions may be less accurate. To illustrate this fact, we repeated the CPD protocol using Gasteiger-Marsili charges (without pi contribution^50^, as implemented in Open Babel^51^) as non-native charges, and successfully folded (AAQAA)3 starting from its linear structure (S3 Fig). The key difference between the force field-native charges and EEM charges is related to conformational dependence (e.g., side-chain flipping is reflected in EEM charges) and inter-residue charge transfer (i.e., residues will have non-integer charge according to EEM). Despite the fact that Gasteiger-Marsili charges provide a more limited description of the electron density distribution, they can be used to perturb an MD force field that was optimized to work with other types of charges.

Perturbing the atomic charges results in perturbing an isolated component of the forces, which is, in essence, similar to effect of enhanced MD strategies such as Hamiltonian replica exchange^52^. Another similarity is that multiple simulations are run using different energy functions, especially if the non-native atomic charges depend on the molecular conformation (i.e., each perturbed segment will use different forces). However, unlike replica-exchange MD, CPD uses sequential rather than parallel simulations.

### Role of the force field

The use of a certain force field can be critical for obtaining correct results. For example, ff03, which is the force field used in our calculations, is known to overstabilize helices^53^. To determine whether CPD can help expedite folding even when using a force field that does not have such a bias, we have repeated the CPD protocol using ff99sb-star-ildnp^54^, which belongs to the ff99sb family of force fields, known to underestimate the formation of helices^53^. Using the CPD protocol with ff99sb-star-ildnp, we successfully folded (AAQAA)3 starting from its linear structure (S4 Fig). Importantly, the final folded structure obtained using CPD with ff99sb-star-ildnp is less helical than that obtained using ff03, and more similar to the reference structure (backbone RMSD, 1.21 Å for ff99sb-star-ildnp vs. 1.98 Å for ff03) obtained by Beauchamp et al.^47^, who also used ff99sb-ildn force fields with side chain and backbone torsion modifications (ff99sb-ildn-phi and ff99sb-ildn-NMR, respectively). For comparison, classical MD using the same force field did not achieve folding within the same simulation time (S4 Fig).

Similarly, whether or not non-helical structures can be folded using CPD depends mostly on the force field itself. For example, MD-based studies reported that the force field OPLS-AA can be used to fold the tryptophan zipper (trpzip), a peptide motif that adopts beta-hairpin conformation^55-57^. To determine whether CPD can help expedite the folding of not only helical but also beta-hairpin peptides, we applied the CPD protocol to fold trpzip (PDB ID: 1LE0) starting from the extended structure. For this purpose, we used the OPLS-AA force field. The results (S4 Fig) confirmed that CPD can indeed expedite folding of beta-hairpin peptides provided that the force field is capable of stabilizing such secondary structure elements. Taken together, these observations indicate that the force field is the main determinant of folding effectiveness, whereas charge perturbation is a determinant of folding efficiency.

### Advantages of CPD

We designed CPD aiming to expedite MD-based folding simulations without requiring additional computational resources or expert knowledge. Benchmarking revealed that CPD allows folding of medium-length peptides using the same software, hardware, and know-how required for running classical MD simulations, but less computational time (from the linear extended structure in 45 ns of simulation time). Fast-folding peptides are typically examined using at least several hundred ns of simulation time (e.g., by running several replicas, each totaling tens of ns) and often require applying complex techniques to enhance conformational sampling^58^.

One of the few studies that used extended structures as the starting point of the simulation was performed by Mou et al.^30^, who developed and implemented a new version of the AMBER force field, employed a complex equilibration procedure, and performed a simulation consisting of 12 temperature-specific replica runs of 160 ns each and 2 classical runs of 500 ns each (totaling ∼3 μs), followed by an extensive cluster analysis with the aim to fold the Trp-cage and examine the folding dynamics. Their best structure had a backbone RMSD of 1.1 Å relative to the reference structure with PDB ID 1L2Y, which is the same as the reference structure used in our present study. The same authors obtained three low-energy basins (best RMSD between 1 and 4 Å) that correspond well to the folded structures obtained using CPD in our study. Similarly, Kannan and Zacharias^58,59^successfully folded the Trp-cage (best RMSD, ∼2 Å) from the linear structure by employing biasing potential replica-exchange MD (5 replicas × 70 ns = 350 ns of total simulation time). The same authors later showed that the Trp-cage can also fold from the extended structure in 500 ns of classical MD using various force fields, but with poorer results (C-alpha RMSD >3 Å)^38^. For comparison, CPD provided a backbone RMSD of Å after only 45 ns of total simulation time using a standard force field available in any MD program. Moreover, our short, basic simulation at a single temperature was also able to sample characteristic folding features detected in the complex study by Mou et al.^30^, such as the fact that the Trp-cage first adopts a U-shape, then forms the α-helical stretch, and only afterwards forms the 3_10_-helix.

Another study that used extended structures as starting structures was performed by Lee et al.^60^, who successfully folded Trpzip2 (PDBID: 1LE1; best RMSD, 2.3 Å) and (AAQAA)3 by employing a combination of temperature and Hamiltonian replica-exchange MD (16 replicas × 200 ns = 3.2 µs of total simulation time) using ff96 and implicit solvent. Their simulations provide several low-energy basins that correspond well to the folded structures we obtained using only 45 ns of CPD.

Therefore, while the CPD-based description of the folding dynamics is relatively crude and CPD is not meant to replace long simulations with enhanced sampling, CPD represents an inexpensive yet powerful approach for probing folding dynamics and generating relevant three-dimensional conformations of small proteins based only on information regarding the amino acid sequence.

The literature contains a large and heterogeneous body of computational studies on the folding of the three peptides discussed in our paper. Given that folding for such systems often takes place on a μs time-scale, many studies were successful at modelling folding pathways precisely because they achieved such time-scales (e.g., as did Lindorff-Larsen et al.^17^). In this context, we conclude that, since CPD allows to fold medium-sized peptides in under 45 ns of simulation time, it is at least one order of magnitude faster than any currently available alternative based on MD. Moreover, CPD is applicable to any class of molecules and can be incorporated into any simulation setup, regardless of force field, treatment of solvent, and other methodological aspects. However, the exact CPD protocol should be optimized for specific cases (e.g., by varying the length and frequency of the perturbed MD segments, as well as the type of non-native charges). Finally, CPD does not require a new MD implementation and can be immediately adopted in practice with any MD program.

Importantly, the computational requirements for CPD do not differ from those of classical MD, whereas other enhanced MD simulations are more difficult to set up for inexperienced users and typically require above-average computational resources. For example, if a system requires 12 cores (with certain minimal specifications) to run classical MD, the same 12 cores will be sufficient for CPD, whereas at least n×12 cores will be required simultaneously to run replica-exchange MD (where n is the number of replicas), regardless of how much simulation time is covered. The additional CPD step of atomic charge calculation has no effect on the complexity of the calculation, on the required architecture, or on the overall duration of the calculation (i.e., CPU hours). The only determinant of speed is the force field implementation and simulation setup (e.g., all-atom vs. coarse-grained representation, treatment of electrostatics, water model), as well as the available hardware (e.g., using CUDA acceleration on machines with GPU). These aspects will influence the real-time speed of the calculation (ns/day) regardless of the type of conformational sampling used (classical MD, replica-exchange MD, CPD, etc.).

Furthermore, the nature of the speed enhancement due to improved conformational sampling is important. For example, replica-exchange MD not only helps detect and promote relevant conformers but typically covers more simulation time in less real time by increasing CPU time, provided that sufficient computational power is available. On the other hand, CPD helps achieve folding within a short simulation time, which automatically reduces both the CPU time and the real time required for simulations, and moreover does not require additional computational resources. To summarize, CPD is accessible to any user who can run classical MD.

### Limitations

Several limitations should be considered. First, CPD is limited in its description of the folding dynamics. For example, although folding close to a biologically active conformation can be achieved, such simulations do not necessarily provide the natural folding pathway. Additional limitations are related to the force field itself, which may induce bias towards certain arrangements^47,61^. Moreover, different force fields integrate atomic charge parameters differently, and therefore their sensitivity to perturbation may also differ. Further study is warranted to develop force field-specific CPD protocols that provide efficient folding of small proteins by taking advantage of the particular strengths and weaknesses of each force field, especially in the context of a certain combination of force field, water model, and treatment of electrostatic interactions. These aspects are particularly important when studying molecules with reduced secondary structure content even in the folded state.

It should be noted that, while CPD is a fast alternative to other MD-based techniques, it is more time demanding than fundamentally different approaches to folding, such as those based on Hidden Markov Models. The web server PEP-FOLD is a great example of a widely available tool for rapid prediction of peptide structure starting only from sequence information^62^. On the other hand, non-MD folding approaches provide only the folded structure, while CPD also produces a free-energy profile where transitions of interest can be further studied; moreover, CPD can be used as a conformer generation tool in the computational study of peptides via chemoinformatics or molecular simulation techniques; finally, CPD caters to a wider MD community because it uses common force fields and samples conformations compatible with such force fields, allowing integration with MD pipelines.

## Conclusion

In MD, the behavior of the studied molecule is simulated under the concerted action of a set of forces described using specific mathematical functions with optimized parameters. Using non-native parameters effectively perturbs the MD force field. We showed that this characteristic can be exploited to help expedite folding simulations. In particular, we confirmed that perturbing the electrostatic component of the MD force field can help expedite the folding of medium-length peptides, with successful sampling of important intermediates, using the same software, hardware, and know-how required for running classical (unperturbed) MD simulations. While CPD does not provide an exact description of the natural folding dynamics, it offers certain important advantages over currently available MD techniques in addition to improving sampling: no prior knowledge of the folded or unfolded states is required; there is no need to change the code or settings for classical MD; the perturbation can be achieved using freely available software; regarding computational requirements, CPD is accessible to any user who can run classical MD. CPD can be employed to probe the folding dynamics of known, putative, or planned peptides, as well as to improve sampling in more advanced simulations or to guide further experiments.

## Methods

### Simulation setup

All MD simulations used for benchmarking followed the same protocol and were performed using the GROMACS package^63^. The extended structures used as starting points in each simulation were generated using the AMBER package^64^. The starting structures were placed in a cubic simulation box using the ff03 force field^65^, in such a way that the distance from the solute to any edge of the simulation box was at least 1.5 nm. All bonds were constrained using the linear constraint solver algorithm^66^. Electrostatic and van der Waals interactions were treated via the particle mesh Ewald method^67^, with cubic interpolation and grid spacing of 0.16 nm (or auto-detected when using GPU). The distance for the Coulomb cut-off was 1 nm, and the distance for the Lennard-Jones cut-off was 1 nm (both default values enabling calculations on GPU). The temperature was maintained at 300 K using the v-rescale thermostat^68^with a time constant of 0.1 ps, and Parrinello-Rahman pressure coupling was used^69^, with a time constant of 2.0 ps. The starting structures were energy minimized using steepest decent and equilibrated for 1 ns.

The Leap frog algorithm was used for integrating Newton’s equation of motion, with a time step of 2 fs. In CPD, stage I consisted of 50 iterations, each made up of 400 ps of classical MD plus 100 ps of perturbed MD, giving a total simulation time of 25 ns. Stage II consisted of 20 ns of classical MD. Fully classical MD simulations were run for 100 ns, using only charges native to the force field. Snapshots were taken every 4 ps. All simulations were run in triplicate, with different starting velocities for the atoms. Secondary structure content was evaluated using DSSP^70^in GROMACS. In each iteration of stage I, the classical MD segment was started from the snapshot with the highest content of secondary structure sampled during the preceding segment of perturbed MD; if no suitable structure could be identified (i.e., no residues were involved in a secondary structure element), the last snapshot of the perturbed MD segment was used as a starting frame for the subsequent segment of classical MD. Before starting perturbed MD, the native charges from the topology file were replaced with non-native charges, which were computed using the Electronegativity Equalization Method^26^implemented in ACC^71^. In the ACC calculation, the total charge was +1 e, −1 e, and 0 e for the Trp-cage, Aβ17-34, and (AAQAA)3, respectively; no solvent was included in the ACC calculation; the parameter set EX-NPA_6-31Gd_PCM was used^27^, and the option Full EEM was chosen.

### Evaluating the folding

The secondary structure elements, RMSD, Rg, and free energy were computed within GROMACS using the options do_dssp, g_rms, g_gyrate, and g_sham, respectively. These steps and settings are included in the pseudocode described in the S1 Appendix. The reference folded structures of the Trp-cage and Aβ17-34 were taken from the Protein Data Bank (PDB IDs 1L2Y and 2MJ1, respectively), and the RMSD values are given for the first models of each NMR ensemble. The reference folded structure of (AAQAA)3, which is not available in the PDB, was taken from the work of Beauchamp et al.^38^. The final folded (lowest-energy) structure from each simulation, together with the reference structures, are included in S1 File.

## Supporting information

structure files

## Acknowledgments

We are grateful to Dr. Sushil Kumar Mishra for useful discussions.

