## Supplementary material for "Charge-perturbation dynamics — a new avenue towards in silico protein folding": Figures S1-S4, Appendix S1

**Supporting Information**

**S1 Fig.** Outcomes of a simulation protocol based on charge-perturbation dynamics (CPD) for the folding of tryptophan-cage mini-protein (Trp-cage), amyloid-β peptide Aβ17-34, and the alanine-based pentadecapeptide (AAQAA)3.

**S2 Fig.** Outcomes of classical molecular dynamics (MD) for the folding of tryptophan-cage mini-protein (Trp-cage), amyloid-β peptide Aβ17-34, and the alanine-based pentadecapeptide (AAQAA)3.

**S3 Fig.** Role of charge perturbation in folding the alanine-based pentadecapeptide (AAQAA)3.

**S4 Fig.** Role of the force field in stabilizing secondary structure elements.

**S1 Appendix.** Pseudocode for implementing the charge-perturbation dynamics (CPD) protocol.

**S1 File.** Zipped file containing the final folded (lowest-energy) structures from each simulation, together with the reference structures (not included in this file; provided separately in .pdb format).

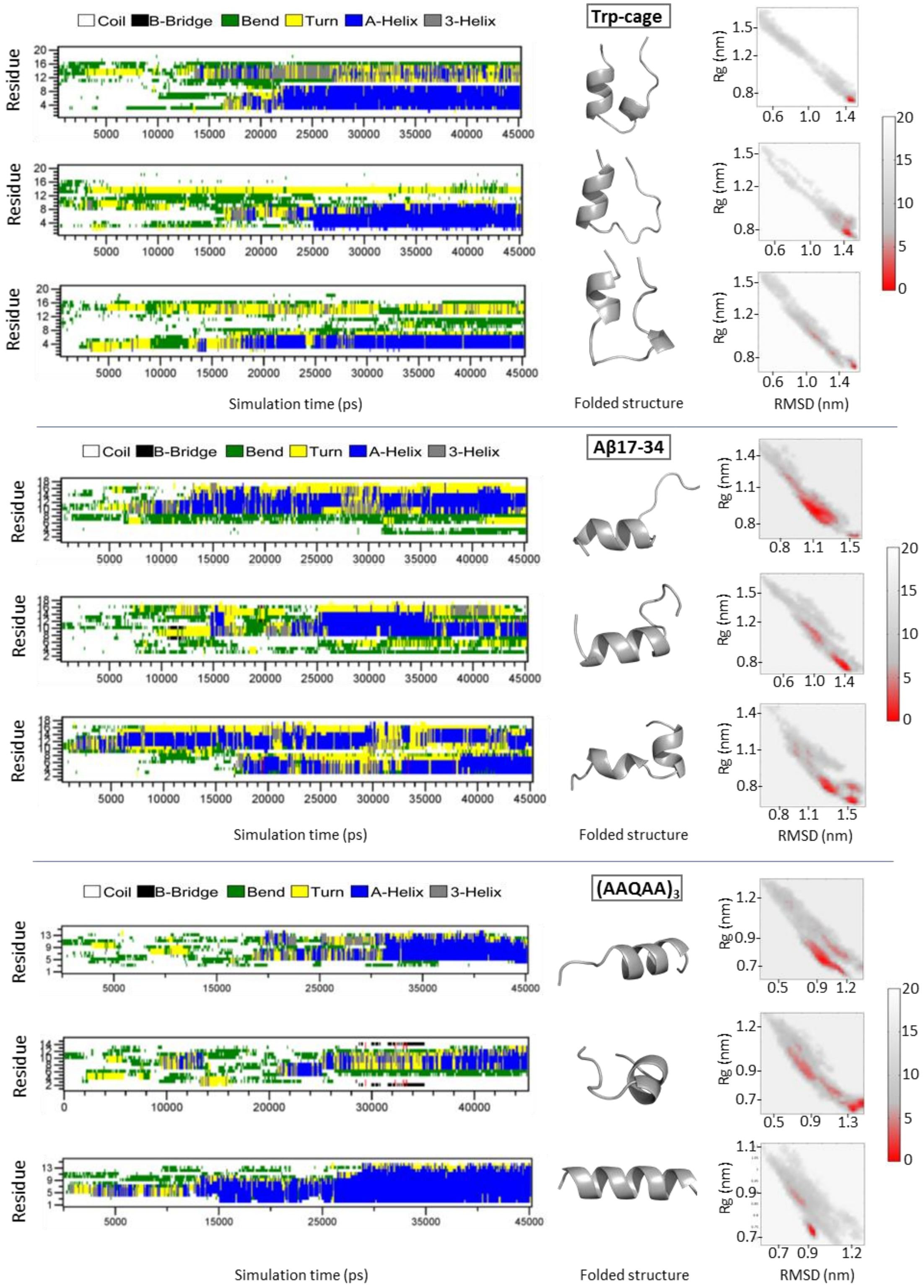


**S1 Fig. Outcomes of a simulation protocol based on charge-perturbation dynamics (CPD) for the folding of tryptophan-cage mini-protein (Trp-cage), amyloid-β peptide Aβ17-34, and the alanine-based pentadecapeptide (AAQAA)$_3$.** The CPD protocol consisted of two stages, amounting to a total of 45 ns of simulation time. All simulations were run in triplicate. Free-energy landscapes and lowest energy structures are provided. Formation of secondary structure elements is also described. Folded structures available in S1 File.

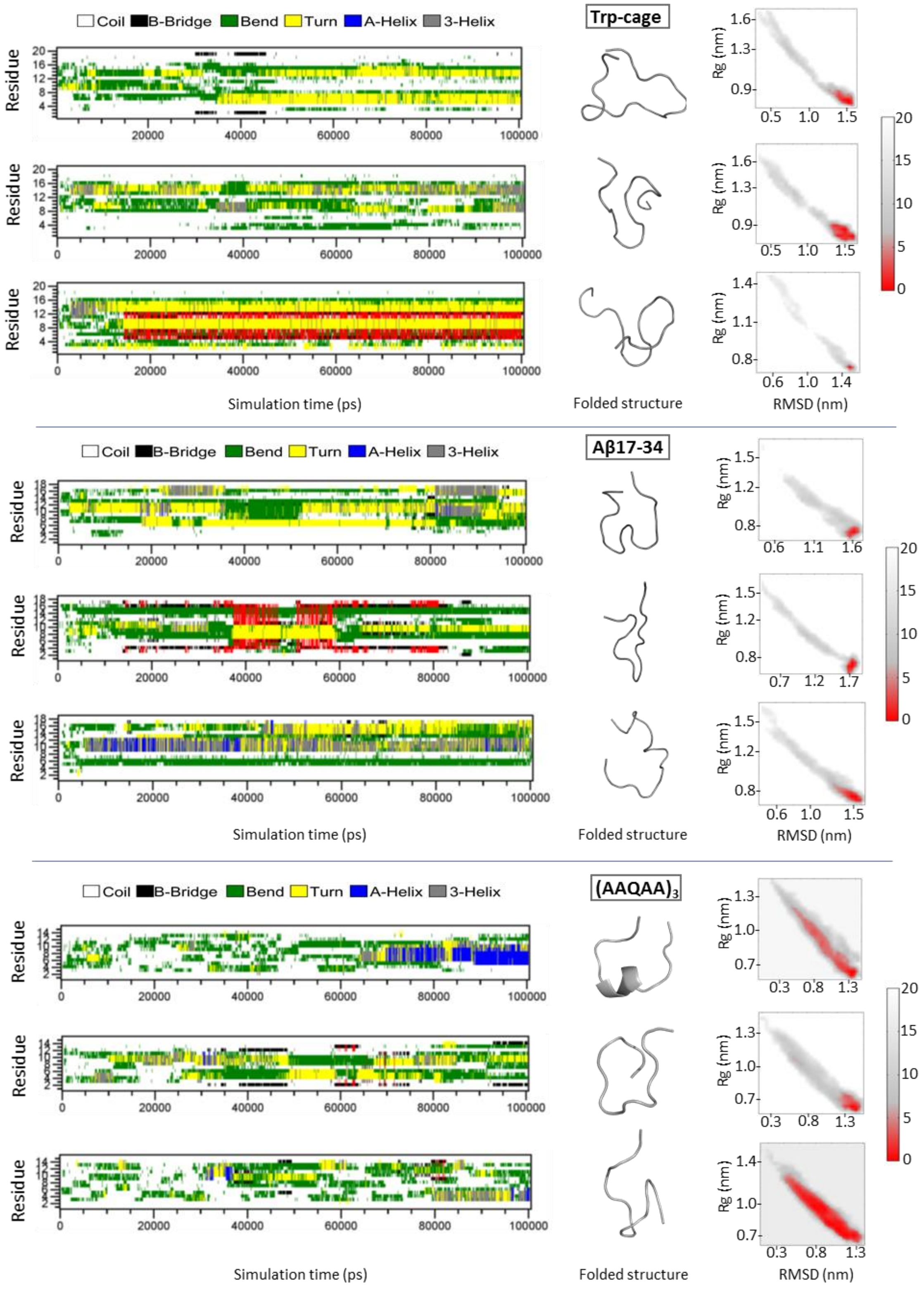


**S2 Fig. Outcomes of classical molecular dynamics (MD) for the folding of tryptophan-cage mini-protein (Trp-cage), amyloid-β peptide Aβ17-34, and the alanine-based pentadecapeptide (AAQAA)3.** A total of 100 ns of classical MD were run. All simulations were run in triplicate. Free-energy landscapes and lowest energy structures are provided. Formation of secondary structure elements is also described. Folded structures available in S1 File.

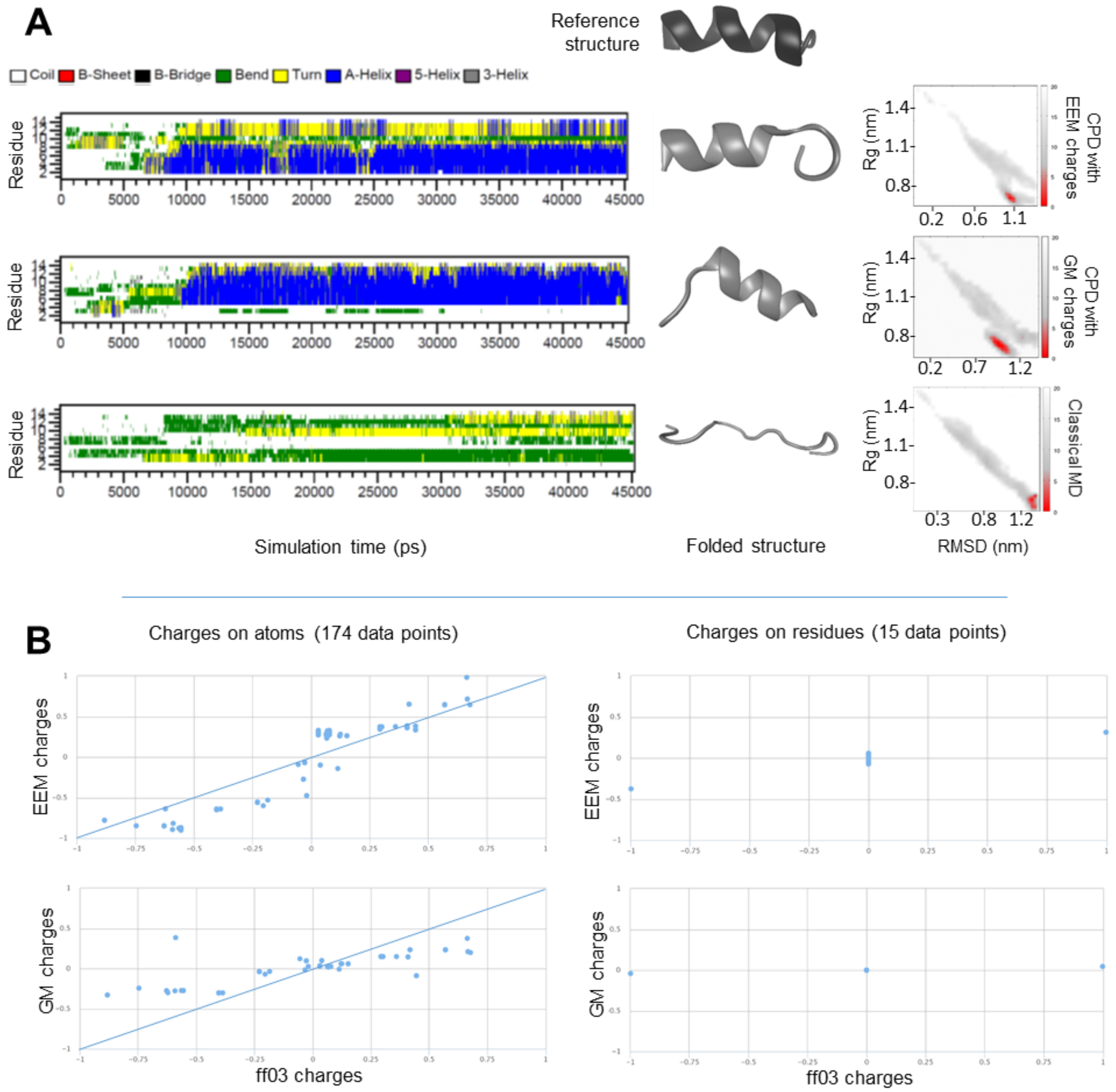


**S3 Fig. Role of charge perturbation in folding the alanine-based pentadecapeptide (AAQAA)3.** (A) The charge-perturbation dynamics (CPD) protocol consisted of stage I (25 ns), where segments of classical MD (400 ps) were intercalated with brief segments of perturbed MD (100 ps), and stage II, where only classical MD was run (20 ns). In CPD, charge perturbation was induced by replacing the atomic charges native to the force field with either EEM or GM charges. For comparison, a classical MD run totaling 45 ns of simulation time is also provided. In both CPD and classical MD, the extended linear structure was used as a starting structure. The simulation setup described in the main text was used, with the following changes: distance between solute and box edges, 1 nm; equilibration, 500 ps; during production, every 500 ps, check if Rg decreased by 5% and, if so, regenerate the simulation box, apply minimization, and equilibrate briefly (100 ps). Formation of secondary structure elements is described. Free-energy landscapes and lowest energy structures are provided, with RMSD against the starting (linear) structure. (B) Comparison of force field native charges (x-axis) against non-native charges (y-axis) for atoms (left column) and residues (right column) for the starting (linear structure). The starting structure is the only common structure in all three simulations described on panel A; however, the starting structure always used the charges native to the force field, since CPD always started with a classical MD segment. Abbreviations: EEM, Electronegativity Equalization Method; GM, Gasteiger-Marsili without pi contribution (i.e., sigma only); MD, molecular dynamics; Rg, radius of gyration; RMSD, root mean square deviation

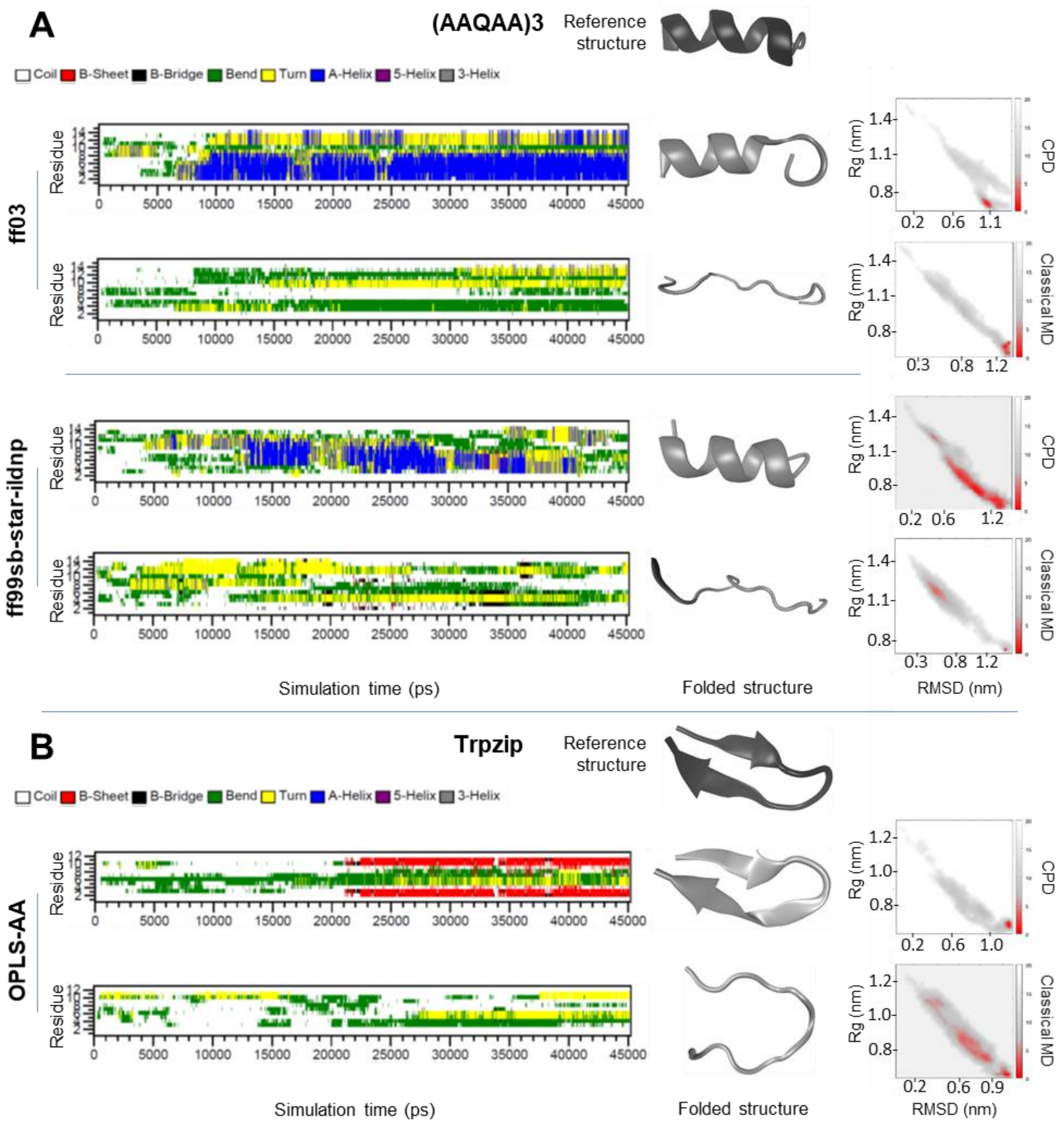


**S4 Fig. Role of the force field in stabilizing secondary structure elements.** The charge-perturbation dynamics (CPD) protocol consisted of stage I (25 ns), where segments of classical MD (400 ps) were intercalated with brief segments of perturbed MD (100 ps), and stage II, where only classical MD was run (20 ns). For comparison, a classical MD run totaling 45 ns of simulation time is also provided for each force field. In both CPD and classical MD, the extended linear structure was used as a starting structure. The simulation setup described in the main text was used, with the following changes: distance between solute and box edges, 1 nm; equilibration, 500 ps; during production, every 500 ps, check if Rg decreased by 5% and, if so, regenerate the simulation box, apply minimization, and equilibrate briefly (100 ps). Formation of secondary structure elements is described. Free-energy landscapes and lowest energy structures are provided, with RMSD against the starting (linear) structure. (A) The alanine-based pentadecapeptide (AAQAA)3, which has high helical content, is folded successfully using CPD with either ff03 (known to emphasize helices) or ff99sb-star-ildnp (from the ff99sb family, known to underemphasize helices). (B) The tryptophan zipper (trpzip), which is a beta hairpin, is folded successfully using CPD with OPLS-AA. Abbreviations: MD, molecular dynamics; Rg, radius of gyration; RMSD, root mean square deviation

**S1 Appendix. Pseudocode for implementing the charge-perturbation dynamics (CPD) protocol.** The pseudocode describes a fully customizable implementation that interfaces with GROMACS, AMBER, and ACC, assuming such programs are accessible. If equivalent programs are installed for molecular dynamics or charge calculation, the corresponding lines in the code need to be changed.

```
//Generate linear structure from sequence using AmberTools
        xleap –f leaprc.FORCE_FIELD
        structure = sequence { N_AMINO_ACID … … C_AMINO_ACID }
//--------------------------------------------------------------------//
//Prepare linear structure for stage I
        //Generate Gromacs input files (.gro, .top)
        echo 1 | gmx pdb2gmx –f structure.pdb –o structure.gro –water spce –p topology.top –i topology.itp –ff
amber03
        //Solvate
        gmx editconf –f structure.gro –o structure.gro –c –d DISTANCE_FROM_STRUCTURE_TO_EDGE –bt
BOX_SHAPE
        gmx solvate –cp structure.gro –cs spc216.gro –o structure.gro –p topology.top
        //Minimize
        gmx grompp –f minimization.mdp –c structure.gro –p topology.top –o run.tpr –po mdout.mdp
        gmx mdrun –v –s run.tpr –o trajectory.trr –c structure.gro
        //Make a backup of the minimized structure for later use
        cp structure.gro initial_structure.gro
        //Equilibrate
        gmx grompp –f equilibration.mdp –c structure.gro –p topology.top –o run.tpr
        gmx mdrun –v –s run.tpr –o trajectory.trr –c structure.gro –cpo checkpoint.cpt
//--------------------------------------------------------------------//
//Set up simulation
        //Prepare Gromacs configuration files
        Create run_classical.mdp (specify number of steps for the classical segment of Stage I)
        Create run_perturbed.mdp (specify number of steps for the perturbed segment of Stage I)
        //Prepare ACC configuration files
        Create ACC_results folder
        Configure charge calculation settings in the ACC configuration file (ACC_setup.json)
//--------------------------------------------------------------------//
//Run simulation – Stage I
ITERATION_COUNT = 0;
//Set the length of Stage I in number of iterations involving one classical and one perturbed segment
while (ITERATION_COUNT < TOTAL_ITERATIONS)
{
        //Run classical simulation segment using Gromacs
                gmx grompp –f run_classical.mdp –c structure.gro –p topology.top –o run.tpr
                gmx mdrun –v –s run.tpr –o trajectory.trr –c structure.gro –cpo checkpoint.cpt
        //Extract the trajectory of the classical run
                echo 1 | gmx trjconv –f trajectory.trr –s run.tpr –o complete_trajectory_s1_temp.pdb
```

```
        //Update the complete_trajectory_s1.pdb file
                cat complete_trajectory_s1_temp.pdb >> complete_trajectory_s1.pdb
        // Make a backup of topology file with native charges
        cp topology.top topology_native.top
        //Calculate non-native charges (using ACC) for the last snapshot of the classical segment
        mono WebChemistry.Charges.Service.exe ACC_results ACC_setup.json
        //In the “topology.top” file, replace native charges by non-native charges
        //Run perturbed simulation segment using Gromacs
        gmx grompp -f run_perturbed.mdp -c structure.gro -p topology.top -o run.tpr
        gmx mdrun -v -s run.tpr -o trajectory.trr -c structure.gro -cpo checkpoint.cpt
        //Prepare for the next iteration
                //Select snapshot with the highest secondary structure content from the perturbed segment
                echo 1|gmx do_dssp -f trajectory.trr -s run.tpr -sc dssp.xvg -dssp_figure.xpm
                //Identify the snapshot with the highest sum of residues involved in beta-sheets, beta-bridges, alpha-helices, 5-helices, or 3 helices
                //Extract the snapshot of interest from the trajectory of the perturbed segment and save it as the starting structure (structure.gro) for the next classical segment
                echo 0|gmx trjconv -f trajectory.trr -s run.tpr -o structure.gro -b BEST_FRAME -e BEST_FRAME
                //Restore the native charges into the topology.top file
                cp topology_native.top topology.top
                //Extract the trajectory of the perturbed run
                        echo 1|gmx trjconv -f trajectory.trr -s run.tpr -o complete_trajectory_s1_temp.pdb
                //Update the trajectory file for stage I
                        cat complete_trajectory_s1_temp.pdb >> complete_trajectory_s1.pdb
                ITERATION_COUNT++
}
//-----------------------------------------------------------------//
//Find the lowest-energy snapshot from Stage 1
        //Calculate RMSD between initial structure and each snapshot in the Stage 1 trajectory
                echo 4 4|gmx rms -f complete_trajectory_s1.pdb -s initial_structure.gro -o rmsd_s1.xvg
        //Calculate radius of gyration for each snapshot in the Stage 1 trajectory
                echo 1|gmx gyrate -f complete_trajectory_s1.pdb -s initial_structure.gro -o gyration_s1.xvg
        //Prepare input file for generating the free energy landscape as a function of RMSD and radius of gyration
                cp rmsd_s1.xvg rmsd_gyration_s1.xvg
                Include the data from the second column of the “gyration_s1.xvg” as a third column in the file
        “rmsd_gyration_s1.xvg”
        //Plot free energy landscape based on RMSD and radius of gyration
                gmx sham -tsham 300.00 -f rmsd_gyration_s1.xvg -ls gibbs_s1.xpm -lsh enthalpy_s1.xpm -lss
        entropy_s1.xpm -bin histogram_index_s1.ndx
        //Extract the frame with lowest energy
                From the “gibbs_s1.xpm” and “histogram_index_s1.ndx” files, identify, among the snapshots that
clustered in the lowest energy group, the snapshot with the highest content of secondary structure. This snapshot
will be used as the starting structure of the stage II simulation (structure_s2.gro).
//-----------------------------------------------------------------//
```

```
//Prepare selected structure for Stage II
	//Solvate
		gmx editconf -f structure_s2.gro -o structure_s2.gro -c -d DISTANCE_FROM_STRUCTURE_TO_EDGE -bt BOX_SHAPE
		gmx solvate -cp structure_s2.gro -cs spc216.gro -o structure_s2.gro -p topology.top
	//Minimize
		gmx grompp -f minimization.mdp -c structure_s2.gro -p topology.top -o run.tpr -po mdout.mdp
		gmx mdrun -v -s run.tpr -o trajectory.trr -c structure_s2.gro
	//Equilibrate
		gmx grompp -f equilibration.mdp -c structure_s2.gro -p topology.top -o run.tpr
		gmx mdrun -v -s run.tpr -o trajectory.trr -c structure_s2.gro -cpo checkpoint.cpt
//-----------------------------------------------------------------//
//Run simulation - Stage II
	gmx grompp -f run_s2.mdp -c structure_s2.gro -p topology.top -o run.tpr
	gmx mdrun -v -s run.tpr -o trajectory.trr -c structure_s2.gro -cpo checkpoint.cpt
		checkpoint.cpt
//-----------------------------------------------------------------//
//Extract the trajectory of the stage II run
	echo 1|gmx trjconv -f trajectory.trr -s run.tpr -o complete_trajectory_s2.pdb
		checkpoint.cpt
//-----------------------------------------------------------------//
//Combine the trajectories of stages I and II
	cp complete_trajectory_s1.pdb complete_trajectory_s1_s2.pdb
	cat complete_trajectory_s2.pdb >> complete_trajectory_s1_s2.pdb
		checkpoint.cpt
//-----------------------------------------------------------------//
//Find the folded structure
	//Calculate RMSD between initial structure and each snapshot in the complete trajectory
		echo 4 4|gmx rms -f complete_trajectory_s1_s2.pdb -s initial_structure.gro -o rmsd_s1_s2.xvg
	//Calculate radius of gyration for each snapshot in the complete trajectory
		echo 1|gmx gyrate -f complete_trajectory_s1_s2.pdb -s initial_structure.gro -o gyration_s1_s2.xvg
	//Prepare input file for generating the free energy landscape as a function of RMSD and radius of gyration
		cp rmsd_s1_s2.xvg rmsd_gyration_s1_s2.xvg
		Include the data from the second column of the "gyration_s1_s2.xvg" as the third column in the file "rmsd_gyration_s1_s2.xvg"
	//Plot free energy landscape based on RMSD and radius of gyration
		gmx sham -tsham 300.00 -f rmsd_gyration_s1_s2.xvg -ls gibbs_s1_s2.xpm -lsh enthalpy_s1_s2.xpm -lss entropy_s1_s2.xpm -bin histogram_index_s1_s2.ndx
	//Extract the snapshot with lowest energy
		From the "gibbs_s1_s2.xpm" and "histogram_index_s1_s2.ndx" files, identify, among the snapshots that clustered in the lowest energy group, the snapshot with the highest content of secondary structure. This is the folded structure.
```
